# Comparison of spatio-temporal dynamics and composition in size-fractionated and unfractionated Northwestern Atlantic microbial communities

**DOI:** 10.1101/2025.04.03.647080

**Authors:** Diana Haider, Jennifer Tolman, Robert G. Beiko, Julie LaRoche

## Abstract

Size fractionation is a widely applied approach to target specific microbial size ranges, differentiating between larger particle-associated, and smaller free-living microorganisms. To characterize its impact on microbial diversity and its comparability to unfractionated samples, we analyzed 16 weekly ocean samples across five depths during a spring bloom. A universal marker was used to characterize prokaryotes, eukaryotes and chloroplasts comparing single (0.2 µm) or sequential (3 µm and 0.2 µm) filtration. We analyzed unfractionated, fractionated (small and large fractions) and de-fractionated samples. The particle-associated fraction defines the most different community from the other fractions, and combining size fractions (de-fractionating) before or after sequencing produces a community that is most similar to unfractionated samples in terms of composition, and richness dynamics with the exception of very rare taxa. Between 75% and 97% of features are shared, but some discrepancies in relative abundances were unresolved, including for some lineages of free-living Proteobacteria like OM43. The richness trends were consistent, and ANCOM detected at most one significantly different feature between fractionated, and de-fractionated samples, highlighting the similarity in community composition and temporal dynamics between the sets.

## 1 Introduction

Communities of microorganisms composed of bacteria, archaea, and unicellular eukaryotes dominate virtually all environments on Earth, from the depths of the ocean to complex terrestrial ecosystems. In the ocean, the heterogeneity of microbes attributable to the range of morphological, physiological, metabolic and behavioral differences grants essential and diverse ecosystem services, such as their contribution to biogeochemical cycling of carbon and nutrients Wong [2015], Falkowski et al. [2004]. Profiling of ocean microbiome diversity is routinely done by sequencing a single target gene, commonly the universally present 16S ribosomal RNA (rRNA) gene for prokaryotes or 18S rRNA gene for eukaryotes, at a given sampling location with protocols that target all microorganisms with size larger than 0.2 µm.

Marker-gene-based profiling of the marine microbiome is carried out by filtering a known quantity of seawater through a filter of a given pore size to collect cells. This is followed by DNA extraction from the particulate material collected on the filter and partial or complete sequencing of universal gene markers like 16S rRNA or 18S rRNA. Size fractionation during the filtration step is a common practice in field sampling that is used to either increase the volume of water filtered through very small pore size (eg. 0.2 µm) or to target specific groups of larger microorganisms (e.g. *>* 3 µm). Size fractionation increases labor and costs associated with marine microbiome research, but provides enhanced recovery of organism diversity with insights on the ecological functioning, trophic dynamics, growth and metabolic rates often associated with various size characteristics Pommier et al. [2008], Massana et al. [2015], Benedetti [2021], Kavagutti et al. [2023]. Fraction-specific taxonomic profiles can differentiate between same-species life cycle stages, free-living and particle-associated prokaryotes, and pico-, nano-, micro-, and meso-plankton Pascoal and Tinta [2023], Tao and Hou [2022], Mestre et al. [2018]. Microbes of different sizes in ocean microbiomes occupy distinct ecological niches, and contribute uniquely to ecosystem services Finlay [2002], Wu et al. [2017], Tao and Hou [2022].

The Bedford Basin is a semi-enclosed 71m deep bay located in the northwestern part of the Halifax Harbour, on the Atlantic coast of Nova Scotia Raes et al. [2022], Robicheau [2022], Robicheau et al. [2023]. The surface waters have limited freshwater input, and a narrow sill connecting it to the open Atlantic Ocean leading to a stratified water column with saltier depths, and fresh waters at the surface Li and Dickie [2001], Rakshit et al. [2023]. Longer day length and warmer temperatures, and ensuing stratification of the water column, trigger spring blooms in the Bedford Basin that are dominated by a few species of phytoplankton that lead to a strong decrease in alpha diversity. While size fractionation during a bloom period may improve the resolution of identified microbial species, the labor and cost-intensive nature of size-fractionation led us to assess the differences between size-fractionated and unfractionated samples.

In this study, we explore the effects of using size fractionation to recover large (*>*3 µm) and small (0.2-3 µm) organisms compared to an unfractionated sample where all particles *>*0.2 µm are recovered on a single filter. We assess how to reconstitute size-fractionated samples for the most accurate comparison with unfractionated samples, and discuss instances when size fractionation is advantageous, for example during a spring bloom. We used sequence reads from a universal primer targeting both 16S and 18S rRNA genes Parada et al. [2016], Walters et al. [2016] in weekly seawater samples collected before, during and after the spring bloom of 2022 in the Bedford Basin as part of a larger sampling effort of the Bedford Basin Monitoring Program.

We tested the effects of de-fractionation using a multi-depth and temporal amplicon dataset from the Bedford Basin generated from 500 mL water samples and amplified with universal primers Parada et al. [2016], Walters et al. [2016]. DNA sequences from fractionated and unfractionated samples were assigned to prokaryotic, chloroplast, and eukaryotic taxonomic groups. We focused on two approaches that provide similar results to unfractionated samples: combining the size fractions in proportion to their DNA concentrations relative to the total DNA of both fractions either before or after sequencing. We show that these samples provided the highest similarity to unfractionated samples, and that size fractionation allows for the description of particle-associated community members which can be masked in unfractionated samples. Both approaches were thus considered effective in comparing water samples that were collected in parallel with and without size fractionation with discrepancies only observed in very rare features.

## 2 Materials and Methods

### 2.1 Sampling and environmental data

For 16 weeks between January and April 2022, weekly water samples were collected using Niskin bottles at depths of 1, 5, 10, 30, and 60m at the Compass Buoy Station in the Bedford Basin, Halifax, Nova Scotia, Canada (44.6936 N, 63.6403 W) with the exception of week 1 30m which was not sampled. 500mL of seawater from each depth was passed either sequentially through 3µm and 0.2µm filters (size-fractionated) or directly onto 0.2µm (unfractionated) 47mm polycarbonate membranes (Millipore) using a peristaltic pump. Filters were flash frozen in liquid nitrogen and maintained at -80°C until eDNA extraction. Nutrients concentrations such as phosphate, silicate, nitrate, and ammonia, alongside chlorophyll a, and temperature data were provided by the Bedford Basin Monitoring Program (https://www.bio.gc.ca/science/monitoring-monitorage/bbmp-pobb/bbmp-pobb-en.php).

### 2.2 Extraction and sequencing

DNA extraction and sequencing followed the protocols described in Robicheau et al., Robicheau [2022]. eDNA was extracted from filtered biomass using the DNeasy Plant Mini kit (Qiagen) with an enhanced lysis procedure. Filters were incubated for five minutes at room temperature with 50 µL of 5mg/mL lysozyme (Fisher BioReagents), then with 45µL of 20mg/mL proteinase K (Fisher BioReagents) and 400µL of Buffer AP1 at 52°C for 1h with continuous shaking in a ThermoMixer (Eppendorf). Subsequent extraction procedures followed the manufacturer’s protocol; eDNA was eluted in 100µL of Buffer AE, and the concentration measured with a NanoDrop 2000c (Thermo Scientific). The V4-V5 region of the 16S rRNA gene, and V4 region of the 18S rRNA were sequenced on an Illumina MiSeq at the Integrated Microbiome Resource (Dalhousie University, Halifax, Nova Scotia, Canada) using the universal primers 515FB-GTGYCAGCMGCCGCGGTAA and 926R-CCGYCAATTYMTTTRAGTTT Parada et al. [2016], Walters et al. [2016].

### 2.3 Data processing

Sequence data was split into 16S rRNA and 18S rRNA amplicons using *bbsplit* (BBMap suite: Bushnell [2014]) to bin reads with their respective target region using SILVA (Quast et al. [2012]) and for the remainder of the analyses the 16S and 18S rRNA regions were processed separately. Unlike the 16S rRNA, the 18S rRNA target region is longer, yielding non-overlapping paired reads. As a consequence, the forward and reverse reads were concatenated following the approach of Needham et al. 2019 (Mcnichol. 18S rRNA data was pre-processed using packages within *bbmap-env* : *bbduk* for trimming, and *fuse* for merging forward and reverse reads Bushnell [2014]. In contrast, 16S reads were trimmed and merged with DADA2 using a trimming setting of 280 nucleotides (forward) and 220 nucleotides (reverse). Both datasets were then denoised for amplicon sequence variant (ASV) using *DADA2* within QIIME2 (v. 2023.5.1) Callahan [2016], Bolyen et al. [2019]. From this stage, the ASV units described will be referred to as features. Chloroplast-associated features were removed from the 16S rRNA and re-classified using *PhytoRef* Decelle et al. [2015] for phytoplankton, and SILVA Quast et al. [2012] was used for the prokaryotes and eukaryotes classification. Finally, we combined the feature data with the metadata collected through the Bedford Basin Monitoring Program Li and Dickie [2001].

### 2.4 De-fractionated sets

To assess the effect of size fractionation on the characterization of the microbial community, we categorized samples into four different size classes: small fraction (S, 0.2-3 µm), large fraction (L, *>*3 µm), unfractionated whole (W, *>*0.2 µm), and de-fractionated small-large (SL, combined S and L). The de-fractionated (SL) fractions was generated by combining S and L sequences weighted by their DNA concentration according to Equation 1. Normalization by the DNA concentration of each fraction compensates for the difference in biomass yield between the large, and small fractions. To calculate the feature profile of SL, we combined the features from each fraction (S or L), and calculated the relative abundance of feature i in sample j (*C*_*ij*_) by normalizing the Observed relative abundances (*O*_*ij*_) with the total DNA concentration of the sample (Equation 1).

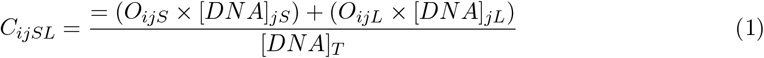

Where *O*_*ijX*_ is the observed relative abundance of feature i in sample j from the size fraction X, *C*_*ijSL*_ is the calculated relative abundance of feature *i* in sample *j* from the size fraction SL, *DNA*_*jX*_ is the DNA concentration of sample *j* from the size fraction X, and *DNA*_*T*_ is the sum of DNA concentration of sample *j* from the combined L and S fractions. While there was four fractions compared across all depths, and weeks, a subset of 18 samples were selected for additional sequencing based on their Aitchison distance between the SL and W fractions (see Supplementary Table 1 for highest and lowest Aitchison distances). These samples were the most dissimilar (largest distance), and most similar (smallest distance) samples. For this pooled (P) fraction, DNA extracted from the small and large filters were pooled at a 1:1 volume ratio before sequencing and was sequenced as a single sample. In contrast, SL was generated by sequencing the small and large fractions separately, and merging them post-sequencing using Equation 1.

### 2.5 Statistical testing

Alpha-diversity comparisons was assessed with the Shannon index and richness counts calculated on unrarefied data. While there is still debate around rarefaction for compositional data McMurdie and Holmes [2014], Willis [2019], Weiss and Lozupone [2017], Schloss [2024], we opted for non-normalized counts because some of our samples (specifically from the large size fraction) have a low library coverage (see Supplementary Figure 5 for rarefaction curves, and Supplementary Figure 6 for DNA concentrations). In this case, rarefaction would cause the loss of a large number of reads. To compare the size fractions by their number of features, we ran an ANOVA with post hoc pairwise t-tests with Holm correction for p-values. The richness was calculated as the total number of unique features per sample, resulting in one richness value per week, depth, and size fraction. The null hypothesis of the t-test is that the mean richness difference between pairs of fractions is zero.

The result of the test is used as an indicator of the richness similarity between the size fractions. Since we have temporal data, we investigated abundance patterns over time through trend analyses, defined as the rate of community richness change over time as either increasing, decreasing, or constant. The trend was characterized through the slope of the linear regression of the richness with time. The time-series analysis was used to assess whether the richness was stable across space, time and size fractions, and to evaluate the trend. To compare the composition between size fractions, we calculated the number of shared features between the size fractions. The sets of features are derived from single fractions, or the union or intersection of the size fractions S, L, W, SL. We calculated weighted proportions of relative abundances to determine the difference between the taxonomic composition of size fractions using Equation 2.

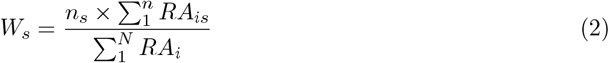

Where *W*_*s*_ is the weighted proportion of a set *s, RA*_*is*_ is the read count of a feature *i* in set *s, n*_*s*_ is the number of features in the set *s*, Nis the total number of features in all sets, and *RA*_*i*_ is the read count of a feature *i*. As a result, the weight of a set is the sum of the read counts of each feature, multiplied by the feature count in a set divided by the total read cound of all features in all sets.

The differential abundance analysis identified individual taxa that significantly differ in relative abundance between groups of size fraction S, L, SL and W. We used the Analysis of Composition of Microbiomes (ANCOM) method from scikit-bio to first normalize the read tables with centered log-ratio (clr) transformations appropriate for compositional data, then compare the relative abundance of each feature pairwise across groups Mandal et al. [2015]. ANCOM generates a W-statistic for each feature which represents the number of times a test showed the feature was significantly different. A high W-statistic suggests a feature is highly differentially abundant between groups, making it an important feature to distinguish between the groups which in this study are the size fractions.

### 2.6 Data availability

Statistical analyses and data visualizations were all generated using python version 3.8.16 and R version 4.3.3. The accession number for the BioProjet in NCBI is PRJNA785606 and all scripts used to analyze and generate the figures are available at https://github.com/dianahaider/size_fractions.

## 3 Results

### 3.1 Environmental variation from January to April

The sixteen weekly samples were taken from January to April 2022, covering surface (depth 1m) water temperatures ranging from 0.7 °C to 6.6 °C. The coldest and warmest temperatures were observed on February 7 and April 27, respectively. The chlorophyll *a* (Chl*a*) fluorescence, which is used as a proxy for phytoplankton biomass, remained below 1.1 µg/L until week 8 (March 1), and thereafter, increased to a peak value of 12.3 µg/L at week 12 (March 12) (Figure 1). The concurrent drastic drawdown of macronutrient concentrations (silicate, nitrate, phosphate, and ammonia), observed at week 12 (March 31) is also characteristic of coastal spring bloom dynamics Robicheau et al. [2023]. Due to the notable changes, we described weeks 1 to 8 (January 1 to March 1), and 9-16 (March 9 to April 27) as pre-bloom and bloom periods, respectively.

**Figure 1:**
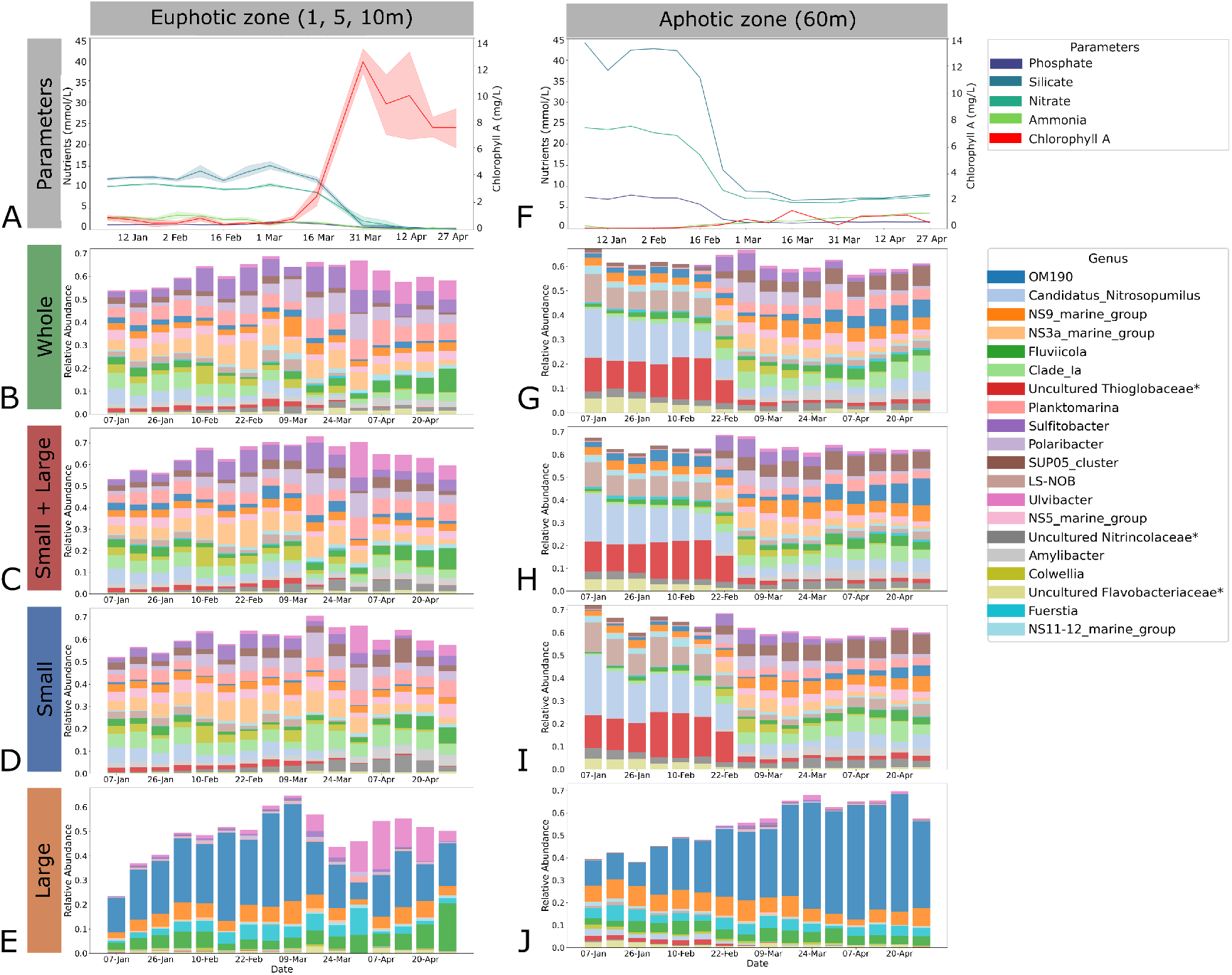
Concentrations of nutrients and chlorophyll a and prokaryotic microbial composition in the unfractionated (whole), de-fractionated (small + large) and fractionated (small, large) sets. Bar plots of the concentrations of nutrients and chlorophyll a in the euphotic (1m, 5m, and 10m) and aphotic zone (60m) measured from January 7 to April 27, 2022. Nutrients including phosphate, silicate, nitrate and ammonia are quantified in mmol/L and chlorophyll a in µg/L. In the euphotic zone, solid lines represent the mean values across depths 1m, 5m, and 10m and the shaded area indicates the 95% confidence intervals. Nutrient concentrations are represented by blue and green hues, and chlorophyll a in red. A secondary y-axis was used for chlorophyll a to account for different scales facilitating the visualization of its relationship with the nutrients. Below the nutrient plots, four taxonomic bar plots show the microbial composition for the whole (B, G), de-fractionated (C, H), small (D, I) and large (E,J) size fractions during the spring bloom depicting the 20 most abundant genera. Some abundant genera were labeled as unclassified or uncultured. To provide more clarity, they were reidentified as uncultured or unclassified, along with their closest available taxonomic rank (e.g., genus, family, or class, depending on availability). These labels are also marked with an asterisk. The 5m depth taxonomy was used for representing the euphotic zone. The taxonomic bar plots for chloroplast and eukaryotes are available in supplementary (Supplementary Figures 3, 4).

In surface waters, we observe the shift from diverse communities to a few plastid ASVs, representative of the dominant phytoplankton species. The phytoplankton community shifted from a dominance of *Teleaulax, Bathycoccus, Thalassiosira* and *Micromonas* during the pre-bloom period to free-living *Pseudopinella* and *Dictyophyceae* or particle-associated *Chaeotoceros* during the bloom. These changes are observable in both the W and SL sets. Bacteroidota such as *Ulvibacter, Fluviicola* (detected in L) and *Polaribacter* increased in relative abundance in response to the bloom (see Supplementary Figure 4 for chloroplast taxonomic bar plots). *OM190*, a large cell from the Planctomycetes phylum classified as the most abundant in L, physically sinks during the bloom as it relatively decreases in surface waters and increases in the aphotic zone. This observation can be made with the unfractionated samples, but the pattern is strikingly obvious within the large fraction (Figure 1E, J). Despite the overlap between detected features in SL and W, the size fractionated samples allow for the distinction between free-living and particle associated members. For example, an uncultured SUP05 clade (*Thioglobaceae*) is detected in both W and SL but the size fractionation highlight its dominance in the small fraction, suggestive of its chemolithoautotrophic role in the depths of the Bedford Basin (Raes et al. [2022]). This is also true for *Sulfitobacter* involved in sulfur cycling, and other heterotrophic bacteria linked to the oceanic carbon cycle such as *Planktomarina* and *Clade_Ia* (SAR11) Marques et al. [2020], Gutiérrez-Barral et al. [2021] (Figure 1). During the intense bloom period, multiple unfractionated samples fail to amplify any eukaryotes but reconstituting the large and small provided a better characterisation of the eukaryotes (see Supplementary Figure 3 for eukaryote bar plots). A eukaryote-specific primer is estimated to provide better eukaryotic taxonomic profiles. Overall, SL and W have highly close to taxonomic profiles for abundant prokaryotic and chloroplast genus, but size fractionation provides better resolution for eukaryotes, specifically during the bloom, both in the surface, and deep waters and allow for differentiation of dominant by size, providing insights into function (Supplementary Figure 3).

### 3.2 Small prokaryotes dominate sample richness

Prokaryotes represent 86.5%, chloroplast 12.7%, and eukaryotes 0.8% of the entire sequence library, i.e. all size fractions combined (Table 1). Across all size fractions, prokaryotes were preferably sequenced, and SL consistently had the highest raw richness. The average richness characterized by the number of unique features observed per sample is smallest for microbial eukaryotes (25 ± 15 ASVs / sample), slightly higher for chloroplast (33±14 ASVs / sample), and highest for prokaryotes (366±132 ASVs / sample) independent of the size fraction (Figure 2). This difference in library size is partly due to the primer bias causing eukaryotic sequences to be selected against, the dominance of prokaryotes in pelagic environments, and the contribution of mitochondrial 16S rRNA of eukaryotes to the prokaryotic pool Parada et al. [2016], McNichol et al. [2021]. Paired t-test at each depth revealed no significant difference between the raw richness, or Shannon diversity among S, L and W size fractions for chloroplasts (*R*_*S*_=31, *R*_*L*_=35, *R*_*W*_ =31) and eukaryotes (*R*_*S*_=19, *R*_*L*_=28, *R*_*W*_ =26), but significant differences were observed when compared with with SL (*R*_*SL*_ = 49 for chloroplast, *R*_*SL*_ = 37 for eukaryotes) (see Supplementary Table 2 for all pairwise comparisons). Prokaryotes from the small (S) size fractions are significantly more diverse than the large (L) size fractions (*R*_*S*_=416, *R*_*L*_=247, *R*_*W*_ =378, *R*_*SL*_=514). For all communities, the reconstructed (SL) size fraction had the highest richness suggesting size fractionation followed by reconstruction, captures more microbial features in comparison to unfractionated (W) samples and may be more appropriate for investigation of rare taxa.

**Table 1:** Read depth and number of features observed in each size fraction for prokaryotes, chloroplasts and eukaryotes Breakdown of the number of features and read depth for each sample type. Prokaryotes and eukaryotes are separated using SILVA as a reference, and chloroplasts are extracted from prokaryotic samples identified as “chloroplast”. Due to the overlap of features between small (S) and large (L), the reconstituted small and large fraction (SL) is not the exact sum of features between S and L. The pooled prior to sequencing fraction (P) method was only applied to 18 samples which were selected according to the Aitchison distance of samples reconstituted (SL) against unfractionated (W). The number of features is pooled from all depths, not averaged per depth.

| Community | Size fraction | Number of samples | Read depth | Number of features |
| --- | --- | --- | --- | --- |
| Prokaryotes | L | 79 | 650 987 | 3 048 |
|  | S | 79 | 2 703 318 | 3 223 |
|  | W | 79 | 1 576 180 | 2 935 |
|  | SL | 79 | 4 451 599* | 6 120 |
|  | P | 18 | 1 295 732 | 963 |
| Chloroplasts | L | 79 | 303 259 | 226 |
|  | S | 79 | 246 192 | 184 |
|  | W | 79 | 174 940 | 188 |
|  | SL | 79 | 583 130* | 333 |
|  | P | 18 | 14 863 | 95 |
| Eukaryotes | L | 79 | 20 537 | 346 |
|  | S | 79 | 9362 | 256 |
|  | W | 79 | 12 889 | 284 |
|  | SL | 79 | 22 917* | 588 |
|  | P | 18 | 1 922 | 117 |

**Figure 2:**
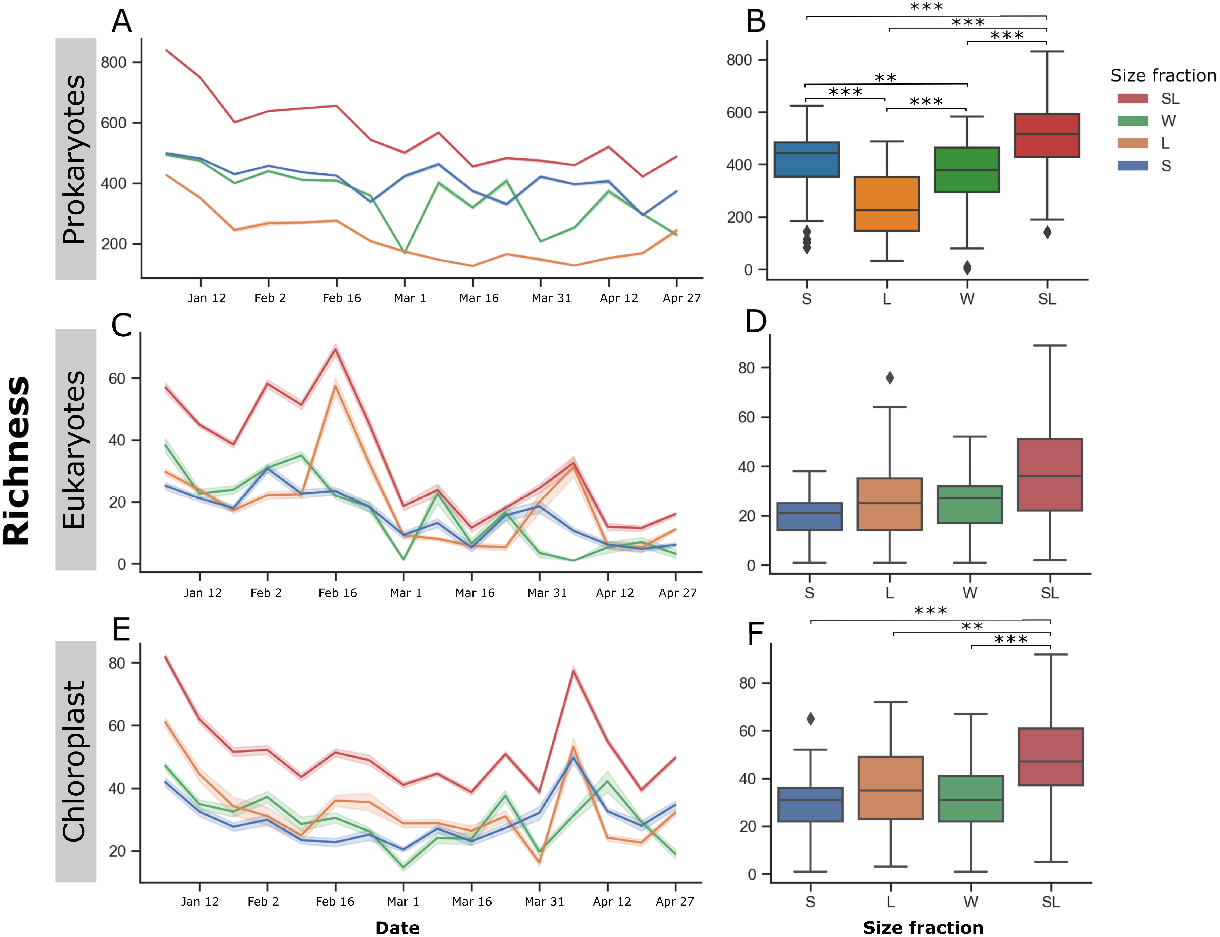
Temporal dynamics of feature counts for prokaryotes, eukaryotes, and chloroplast in the unfractionated, fractionated, and de-fractionated samples. Bar plots of the difference in feature recovery (y-axis) through time (x-axis) by four categories of size fractions for the (a) 16S rRNA, (b) 18S rRNA, and (c) chloroplast communities across 5 depths. The difference is calculated by subtracting the arithmetic mean number of features from the number of observed features for each fraction. As a result, a positive difference indicates that a size fraction has a higher richness in comparison to the average observed for a given week, and a negative difference indicates that the fraction underestimated the richness at a given week in comparison to the other fractions. The boxplots show the richness (y-axis) for each size fraction (x-axis) across all weeks and depths. The median is represented as the line in the middle of the box, and the 25th, and 75th percentiles are the interquartile range (IQR). The whiskers are 1.5 times the IQR, and any outlier is a diamond outside the whiskers. The significance scores denoted by asterisks indicate the p-value from pairwise t-tests (* if p ¡ 0.05, ** if p ¡ 0.01, *** if p ¡ 0.001).

### 3.3 Temporal dynamics varies more by depth than by size fraction

We compared the slope of richness change over time between size fractions to demonstrate alpha-diversity temporal dynamics. The temporal variation remained similar for all fractions, and they were always the same between the pooled (SL) and unfractionated (W) samples (see Supplementary Figure 5 for linear regression scatter plots). Characteristic of spring blooms, the bacterial richness in the upper (1, 5, 10m), and mid (30m) zones decreased from January to April, except for the small size fraction in the deepest (60m) zone which slightly increased in richness (slopeS60m=0.023, Supplementary Figure 5A-E). Chloroplast-associated richness also decreased in the upper zone, and increased at 30 and 60m as the bloom progressed. Eukaryotes from all depths decreased in richness from January to April (Supplementary Figure 5F-J). The slope and trends between S and L varied for certain depth and community combinations highlighting the difference in richness between free-living and particle-associated members of the community throughout the spring bloom. For example, particle-associated chloroplasts increased in richness in the aphotic zone but decreased or remained stable along the water column (Supplementary Figure 5K-O). However, SL and W shared the same trend across depths proving their comparability for tracking richness over time (Supplementary figure 5).

### 3.4 Similarity among samples is strongly associated with size fractionation and bloom time

Beta diversity analyses supported by PERMANOVA testing confirmed the similar community composition of bacterial S, W, and SL, whereas L formed its own cluster. Pairwise PERMANOVA tests confirmed that the three fractions S, W, and SL have minimal variance between each other through large p-values across the five depths, and small *R*^2^ values (Figure 3A, Supplementary Table 3 for PERMANOVA values). For eukaryotes and chloroplast, size fractionation did not result in a clear separation between size fractions, rather they formed temporal clusters from the pre-bloom, to the bloom period (Figure 3, Supplementary Table 3). Our results support that fractionation provides a different large fraction, but that the reconstituted samples are highly similar to the unfractionated samples, and the communities before, and during the bloom are different.

**Figure 3:**
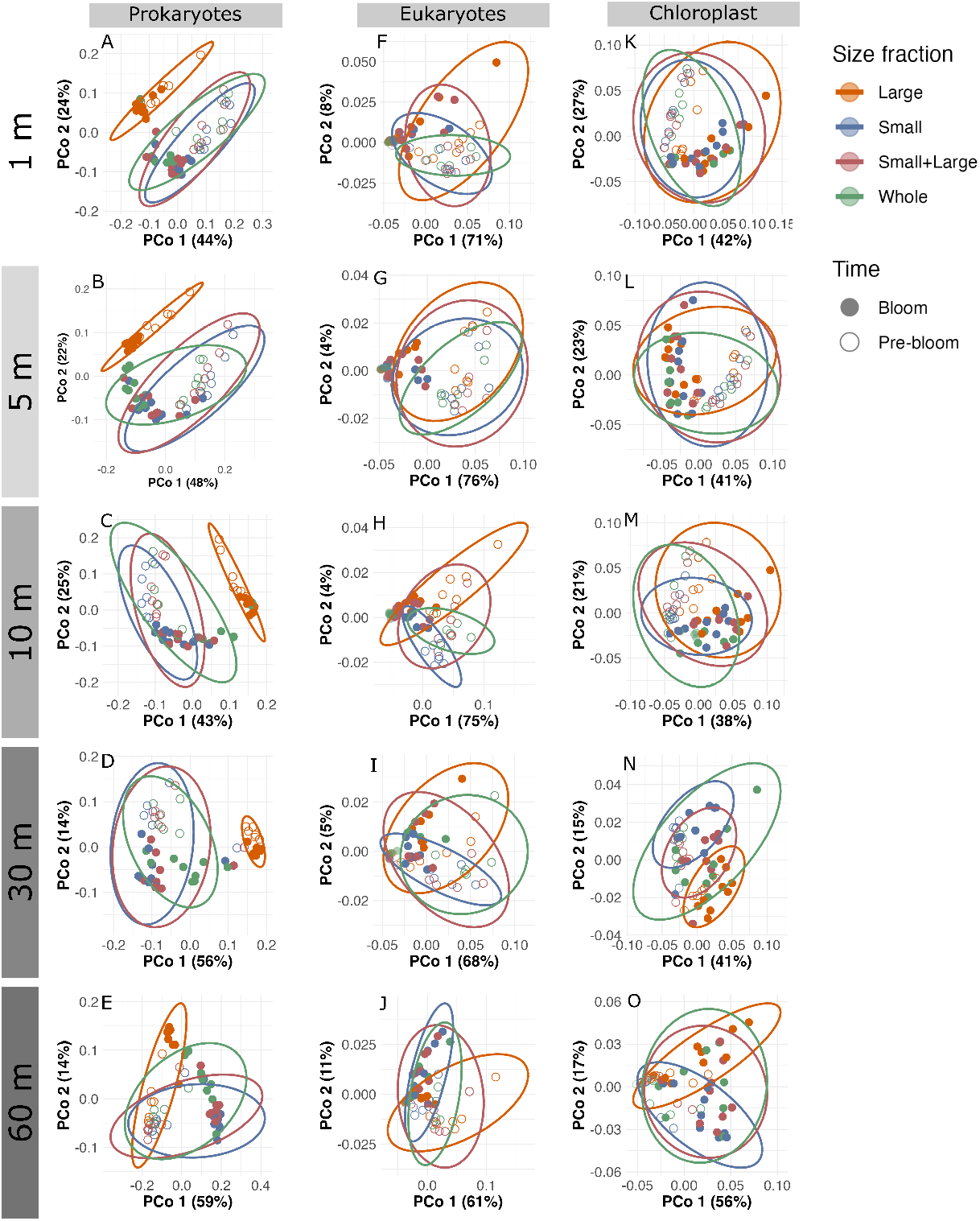
Temporal dynamic of feature counts for prokaryotes, eukaryotes, and chloroplast in the unfractionated, fractionated, and de-fractionated samples. Aitchison distances are displayed in scatter plots for 16S rRNA (left column), chloroplast (middle column) and 18S rRNA (right column). In each PCoA, one point represents a single sample, and the point fill represents the time. The pre-bloom period (open dot) is from weeks 1 to 8, representing the period from January 7 to February 22, and the bloom period (filled dot) is from March 1 to April 27. The colours represent the size fraction, and confidence ellipses were drawn around each fraction in its respective colour where the small is in blue, large in orange, whole in green, and de-fractionated fraction is red.

### 3.5 Most abundant features are shared between size fractions

The majority of abundant features are shared between S, SL and W, and only features that contribute minimally to the overall composition are unshared. Our results show that 70%, 54% and 68% of features are unique to individual size fractions for prokaryotes, chloroplasts and eukaryotes respectively (Figure 4 D, E, F). However, these unique features are very rare (see Supplementary Figure 6 for relative abundance distribution plots of unshared features) and account for less than 3% of the total abundance for prokaryotes and chloroplasts, and less than 25% of microbial eukaryotes (4 A, B, C). When weighted by abundance, we observe that most of the abundant taxonomic classes such as Alphaproteobacteria, Bacteroides, Gammaproteobacteria, Teleaulax, Dinophyceae, Syndiniales and Thalassiosira are shared in all size fractions.

**Figure 4:**
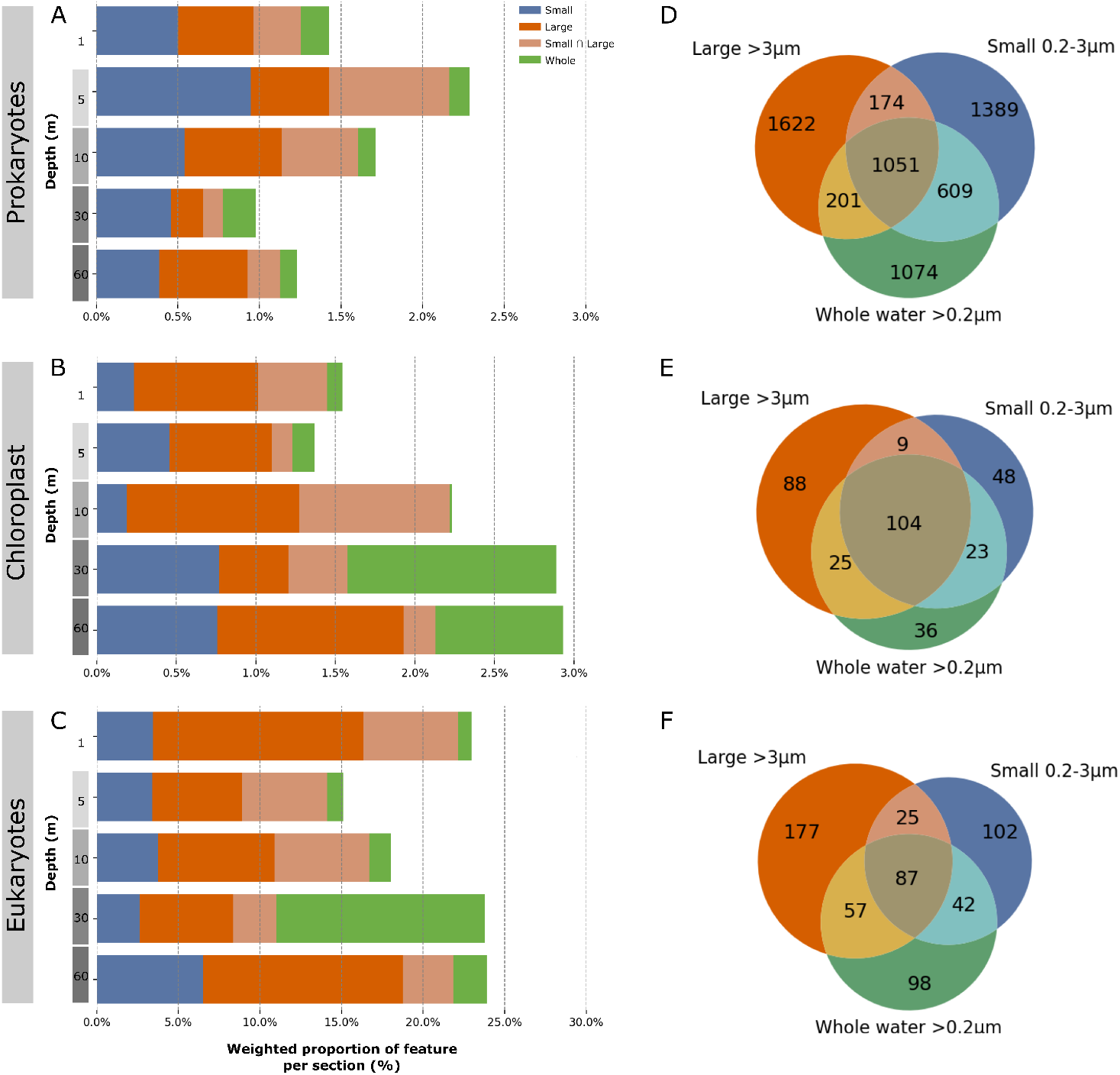
Multi-depth features overlap between unfractionated, fractionated and defractionated prokaryotic, eukaryotic and chloroplast samples. Weighted (a,b,c) and unweighted (d,e,f) proportions of unique features for prokaryotes (a, d), chloroplasts (b, e), and 18S rRNA (c, f). The x axis of the horizontal bar plots is truncated to improve the resolution of the weighted proportion unique to each set. Venn diagrams of the number of shared features between sets of size fractions for d 16S rRNA, e chloroplasts, and f 18S rRNA amplicons. The colours are shared between the right and left plots where the small size fraction is in blue, the large size fraction in orange, the shared ASVs between size and large fraction in light orange, and the whole fraction in green. The brown section (*W* ∩ (*S* ∪ *L*)) represents the features that are commonly found in at least one of the size fractions, and W.

### 3.6 Dominant genera show temporal stability between fractions

Each sample had a most-abundant genus we defined as “dominant”. A total of 16 distinct dominant genera were identified in the prokaryotic communities, 10 in the chloroplast communities and 46 in the eukaryotic communities (Figure 5). These patterns confirm that eukaryotes amplified with the universal primer have greater spatio-temporal variability in taxonomic composition and fractions may be more difficult to reconcile (Figure 5). In contrast, bacterial W, SL, and S shared the same dominant taxa 87% to 100% of the time, and there is less variability in the dominant genera in the pre-bloom period for prokaryotes, eukaryotes and cyanobacteria (Figure 5). We show that W, SL and S have compatible patterns of dominant genera, but there is more variation during the spring bloom, and for the eukaryotes specifically.

**Figure 5:**
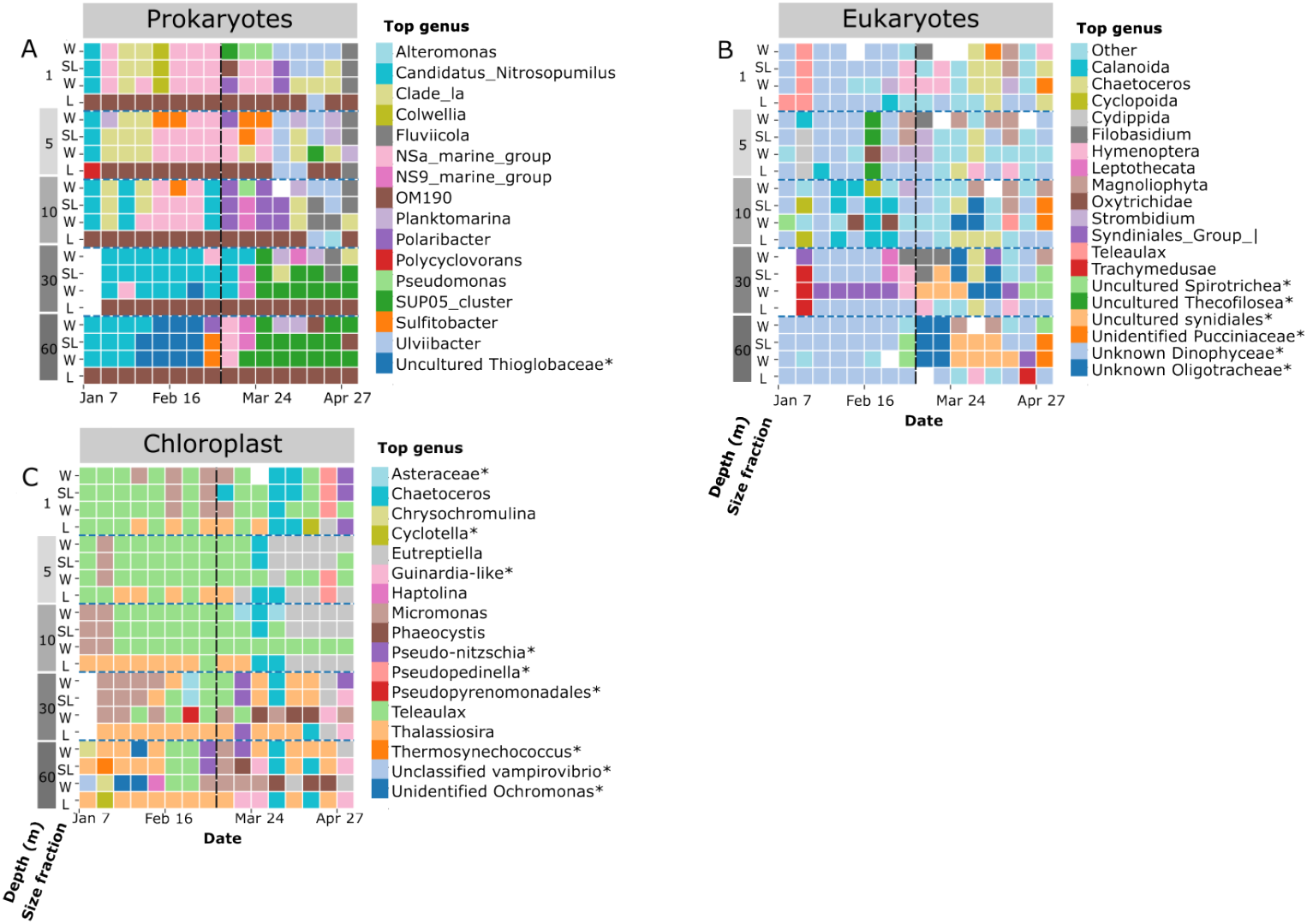
Depth profile of dominant genera in unfractionated, fractionated, and de-fractionated samples during a spring bloom. Dominant genus in the microbial community over a 16-week timespan across depths 1 to 60 m, and four size fractions for a 16S rRNA, b 18S rRNA and c Cyanobacteria amplicons. The 19 most-common 18S rRNA genera are shown, with the remaining 27 grouped into the “Other” category. They include Prymnesiales, Thalassiosira, Ploimida, Cydippida, Thecofilosea uncultured, Unidentified Eukaryota, Cyclopoida, MAST-2, Syndiniales, Mitochondria, Choreotrichia uncultured, Amoebophrya, Pelagostrobilidium, Oxytrichidae, Spirotontonia, MAST-1A, Cryptocaryon, Caecitellus, Capitellida, Helotiales, Dothideomycetes, Leptothecata, Pucciniales, Prymnesiophyceae, Haplozoon, MAST-7A and Pithites. White boxes indicate absence of data, or samples with zero features. The horizontal dotted lines separate depths, and the vertical dotted line separates the pre-bloom and bloom period. Some abundant genera were labeled as unclassified or uncultured. To provide more clarity, they were reidentified as uncultured or unclassified, along with their closest available taxonomic rank (e.g., genus, family, or class, depending on availability).

### 3.7 Important features for size fraction distinction

To further assess differences in relative abundance patterns between size fractions, ANCOM and pairwise ANCOM tests confirmed that minimal features are detected as significantly different between pairs of size fractions. In contrast, when ANCOM on all fractions (multiple testing), there are more different features detected, probably driven by the largely different L fraction. Consistent with the rest of our results, the prokaryote large size fraction had on average two to three orders of magnitude more differentially abundant features in comparison to chloroplast, or 18S rRNA at any depth (Supplementary Table 2 for all pairwise p-values). The pairwise testing between W, SL, and S revealed between zero and seven different features (Supplementary Table 2). Particularly SL, and W had only 2 significantly different features, one detected at 1m, and one at 60m. *Comamonadaceae* a large freshwater bacteria from the surface, and *Moritella* are both entirely absent from W, but detected at *>*0.002 relative abundances in the large fraction, hence their presence in SL. This suggests that while there are differences in the community composition, the variations observed are within acceptable and small ranges which suggests comparability between the fractionation methods for analyzing microbial composition.

### 3.8 Pooled prior to sequencing

To assess whether pooling size fractions before sequencing enhances the comparability to W, we selected 18 samples based on their Aitchinson distance to W (Supplementary Table 1) to filter sequentially through 3 µm and 0.2 µm filters. We picked samples with poor and good representation to assess how pooling prior to sequencing might influence each. The resulting S and L fractions were pooled (P) on a 1:1 DNA ratio. Consequently, P is a size-fractionated sample that was sequenced and processed as a single unit, while SL consists of pooled sequences from separately sequenced S, and L units. In essence, P underwent the same filtration process as S and L but was pooled before sequencing, resulting in a single sequencing step. P were not more similar to W than SL samples (Figure 6). The SL and P sets both showed a high degree of similarity to W, with no statistical difference seen in any of the communities examined (Figure 6). These results suggest that size fractionated samples can be merged prior to sequencing to save on sequencing costs and yield similar results.

**Figure 6:**
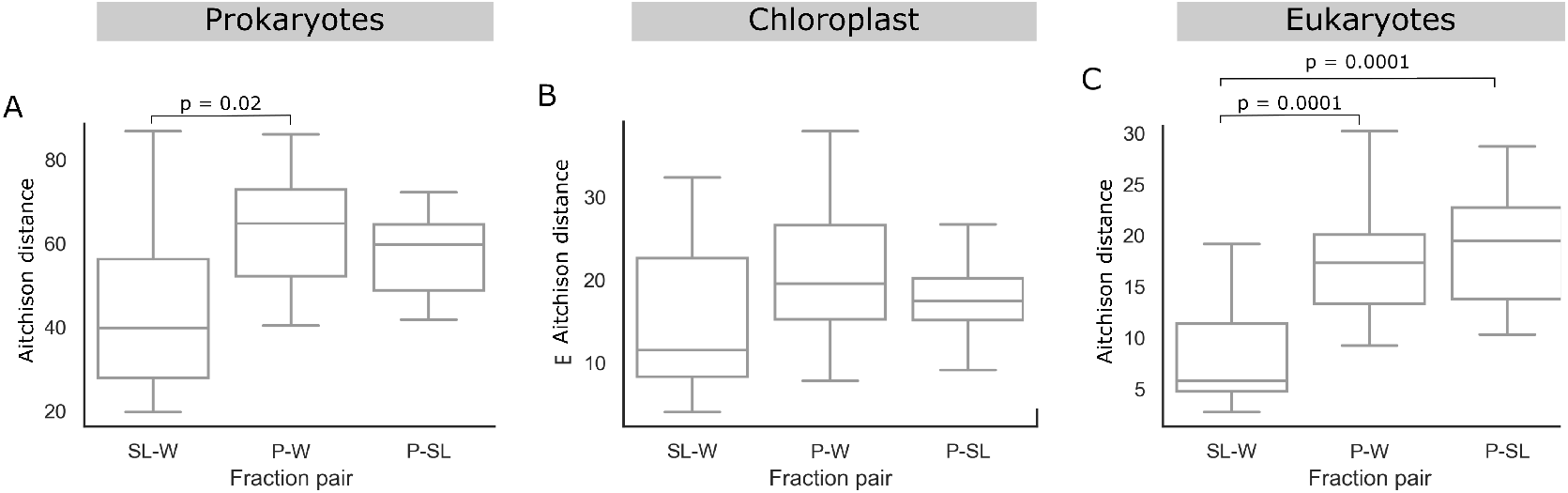
Aitchison distance between 18 pairs of samples that either were unfractionated (W), fractionated and pooled prior to sequencing (P) or fractionated and pooled after sequencing (SL) Boxplot of the Aitchison distance between samples which were processed using different size fractionation protocols comparing the impact of pooling time (pre- or post-sequencing). The boxplot shows the distribution of distances with smaller values indicating greater similarity. Post-hoc t-tests were conducted to assess whether the difference observed between the groups is significant, and significant p-values (¡0.05) were reported on the plots.

## 4 Discussion

In this study, we investigated the impact of size fractionation on the composition and temporal dynamics of the Bedford Basin microbial communities. Our data spans five depths over 16 weeks and captured prokaryotic, chloroplast and eukaryotic communities with a universal primer. Our objective was to assess the similarities between microbial communities inferred from different filtration treatments and to determine how size fractions derived from small, or large pore sizes align with unfractionated samples and whether pooling fractions before or after sequencing impacts the similarity to unfractionated samples.

We found that dominant microbial features are shared across the size fractionated and unfractionaed sets(W). The de-fractionated set where we combined S and L was very similar to W in terms of richness trends, dominant taxa and overall community composition suggesting that size fractionation does not significantly impact abundant community composition. Unsurprisingly, the large size fraction was the most dissimilar to the unfractionated samples, suggesting that particle-associated taxa have distinct community compositions that are highlighted only through size fractionation. However, pooling, either before or after sequencing of the small and large fractions yields communities comparable to those of unfractionated samples in terms of richness, beta diversity patterns, and dominant taxa. Some taxonomic groups such as *OM 190* and *Flavobacteriaceae* exhibited different feature-specific time-series highlighting how size fractionation can capture different changes in relative abundances which are sometimes inapparent or less obvious in unfractionated samples (Table 1).

### 4.1 Environmental dynamics and phytoplankton blooms

Spring phytoplankton blooms typically form in late winter to early spring and are characterized by an increase in sunlight availability and temperature which stratifies the environment (Longhurst [1995], Li et al. [2006]). The upper euphotic zone receives the most sunlight with decreasing light penetration as a function of depth. Phytoplankton species take advantage of these conditions for growth which leads to increased Chlorophyll *a* concentrations, and a rapid utilization and depletion of nutrients (Dai [2023], Needham (2016)). Our data reflects these seasonal changes with W and SL showing high taxonomic similarity during the bloom period but size fractionation highlighting distinct patterns of relative abundance changes between free-living and particled-associated microbes. Despite providing similar dominant community composition, size fractionation is beneficial in low diversity and high biomass conditions, and for capturing rare members of microbial communities. The increased coverage may improve the resolution of rare members in low diversity conditions, and revealing size-specific trends.

In our bacterial and chloroplast communities, the unfractionated (W) taxonomic profile of the 20 most abundant genus through the bloom is highly similar to that of the de-fractionated (SL), for both the euphotic and aphotic zones. However, the communities are more similar between SL and W in their respective relative proportions in the diverse pre-bloom community than during the bloom. This suggests that when diversity is low, size fractionation allows for a better description of the community via increased sequencing depth in comparison to W samples which are often dominated by a few species. By increasing the coverage through two rounds of sequencing, one for each fraction, the chances of sequencing rarer features increases.

### 4.2 Impact of size fractionation on inferred community composition

Our results show that fractionationation using different pore size filters leads to different community compositions, specifically for particle-associated members of the community. However, abundant taxa are shared between the fractions, suggesting the differences are driven by rare features, particularly those with relative abundances ranging from 0 to 0.004 for prokaryotes, and 0 to 0.2 for eukaryotes and chloroplasts (Supplementary Figure 6). SL has the highest richness, which is likely due to the higher sequencing coverage since SL represents two rather than one sample fraction. This is linked to an important methodological consideration that influences richness estimates and comparisons throughout our analysis of samples with different sequencing depths. We reported the number of features without rarefaction because subsampling leads to data loss. As a result, we expect samples with more sequencing depths to have higher richness counts. If samples were rarefied, the richness of the pooled samples would be more similar with the diversity captured in W (Supplementary Figure 1). The particle-associated community consistently had lower richness in comparison to their free-living counterpart.

We used log-ratio transformations Gloor et al. [2017] as an alternative to rarefaction. The highest DNA concentrations were always from the unfractionated samples, and highest starting April 12 (weeks 14, 15, 16), while the lowest concentrations were consistently observed in the large fraction (Supplementary Figure 2). Microbiome data is compositional, and dominant features make up a large portion of the sequencing depth and may consequently mask temporal trends of rarer features. Previous studies showed that filtering out rare features, including low abundance within and between samples can improve reproducibility, and comparability results Schloss [2018], Cao et al. [2021]. However, we retained rare features to provide a complete comparison, as rare features can be important to characterize. In separating the microbial community by size, we enhance the detection of rare features by reducing the effect of dominant species, and obtaining a higher richness in comparison to unfractionated samples. We demonstrate that abundant patterns are well aligned between W and SL but that following the temporal patterns of specific features is challenging and incorrect since some species are misrepresented in S and L McLaren et al. [2019] (Supplementary Figure 7). Therefore, while abundance patterns are similar between SL and W, certain features time-series and the sample absolute richness are different.

### 4.3 Ecological relevance of size fractionation

According to our result, we suggest size fractionation is more beneficial during phytoplankton bloom periods where diversity is lower, and size fractionation increases the detection of rare taxa due to increased sequencing depth. During blooms, few species dominate the community as specific phytoplankton outcompete others and these groups tend to mask less abundant taxa which contribute to the bloom diversity. More specifically, particle-associated microbes are estimated to have higher protease activity and describing size-specific microbial abundances may be more relevant during a spring bloom. In pre-bloom conditions, the communities are more diverse and unfractionated samples capture most of the diversity.

### 4.4 Methodological considerations for size fractionation

While size fractionation can enhance microbial community representation due to a more refined community description by size, our findings suggest that the pooled fraction is very similar in terms of taxonomic composition to the unfractionated samples. While relative abundances differ more than absolute presence absence counts of features, there is evidence that relative abundances from marker gene analyses tend to be inaccurate regardless of the method used. Therefore, if interest is in rare features, specifically divided by their physical size, size fractionation justifies the additional time, and budget allocation. In addition, the fractionation can provide insights into functionality since free-living and particle-associated microorganisms are associated to distinct niches in ocean microbiomes. However, for analyses focused on abundant lineages, we suggest only one filtration step for longitudinal amplicon analysis of microbial communities.

One methodological limitation is the reliance on a single marker gene to profile prokaryotes, eukaryotes and chloroplasts. The primer set we used targets a region that preferentially amplifies the 16S rRNA gene, a marker that is highly effective in capturing prokaryotes Parada et al. [2016]. As a consequence, the eukaryotic diversity is incomplete, and has inconsistent community patterns. Had we employed separate markers, we would have likely achieved a more accurate representation of eukaryotes, however it was beyond the scope of this project. While the use of a single marker simplifies the workflow, it introduces a bias by overrepresenting prokaryotes, and this should be taken into account when interpreting the results of our study.

## 5 Conclusion

We assessed the effect of sequential filtration through large, and small pore size filters, with single-step filtration on microbial community composition and diversity during a spring bloom in a North-western Atlantic microbial community. We show that pooling size-fractionated samples by DNA proportion produces microbial community compositions determined by 16S rRNA amplicons highly similar to those of unfractionated samples which underwent a single filtration step. Our analysis reveals that the small size fraction closely resembles unfractionated samples, and the large size fraction consistently represents a distinct community. However, when small and large samples are pooled, the majority of abundant taxa are shared with the unfractionated samples, indicating that size fractionation does not necessarily provide additional insights into the dominant microbial communities. However, size fractionated samples have higher richness and provide insights into rare taxa that are poorly characterized in unfractionated samples. This enhanced representation of rare taxa is due to the inherent nature of compositional data product by high throughput sequencing, splitting, and separately sequencing samples creates more space to capture rare taxa that are otherwise obscured or undetected in unfractionated samples. We show that pooling fractions prior to or after sequencing does not linearly impact the data and thus yields highly similar results, and therefore is an appropriate method to compare unfractionated with fractionated samples.

## Acknowledgements

This research is part of the large research project of CFREF Ocean Frontier Institute on the dynamic response of microbial communities to changes in water chemistry and water temperature. Funding was also provided by the National Sciences and Engineering Research Council of Canada (NSERC) to Robert Beiko. The authors thank Drs. Erin Bertrand and Alastair Simpson for their helpful conversations and the group of Bedford Basin team for their sampling effort. The Bedford Institute of Oceanography (BIO) and the Department of Fisheries and Oceans (DFO) is acknowledged for assistance in sample collection and for providing physicochemical data.

## 7 Data availability statement

The data that support the findings of this study are openly available in the National Center for Biotechnology Information (NCBI) under BioProject PRJNA785606. The scripts used to generate all the results are available on GitHub: https://github.com/dianahaider/size_fractions.

