## Supplemental information for "Comparison of spatio-temporal dynamics and composition in size-fractionated and unfractionated Northwestern Atlantic microbial communities"

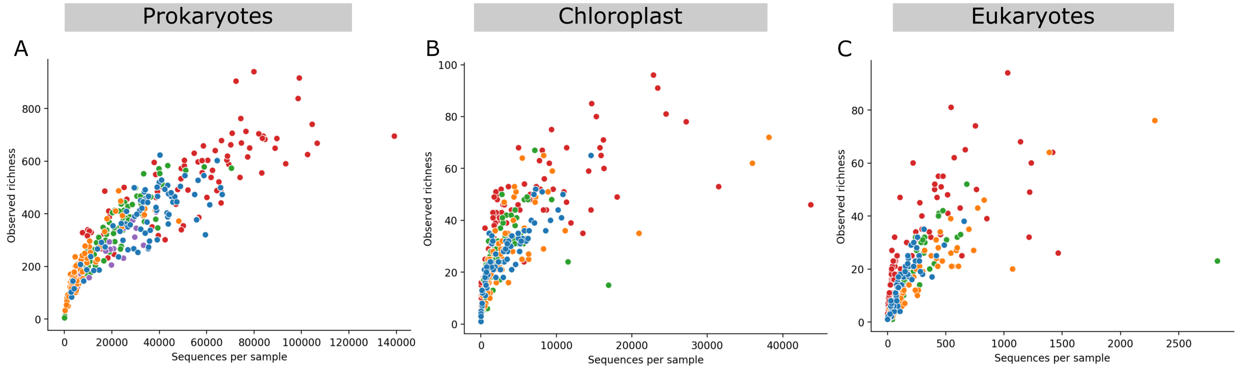


Figure 1: **Rarefaction curves of richness by sequencing depth**
Rarefaction curves illustrating how observed richness increases with sequencing depth, and highlights the problem associated with

**Community (Aitchison distance)**

| | Prokaryote | Eukaryote | Chloroplast |
|---|---|---|---|
| Most similar | **BB22.14A (40)** | BB22.14A (6) | BB22.15C (10) |
| | BB22.12B (42) | **BB22.7D (6)** | BB22.1D (11) |
| | **BB22.15C (48)** | **BB22.12B (7)** | **BB22.5D (11)** |
| | BB22.9A (52) | **BB22.12A (7)** | BB22.3D (11) |
| | **BB22.15B (54)** | BB22.10C (7.33) | **BB22.2D (11)** |
| Most dissimilar | **BB22.10C (107)** | BB22.6C (28) | **BB22.13C (46)** |
| | BB22.13C (103) | **BB22.6D (37)** | **BB22.12C (39)** |
| | **BB22.11E (103)** | BB.4E (27) | BB22.10C (35) |
| | **BB22.9B (101)** | BB22.7C (27) | **BB22.13B (34)** |
| | **BB22.12D (94)** | **BB22. 7E (27)** | **BB22.14E (34)** |

Table 1: **Aitchison distance based identification of most similar and dissimilar samples**
Calculated Aitchison distance between pairs of unfractionated (W) and de-fractionated (SL) samples from the same week, and depth to determine the most similar and most dissimilar samples. The bolded sample entries are those that were selected for re-sequencing as pooled. These samples were pooled in a 1:1 DNA ratio before sequencing to compare whether pooling size-fractionated DNA prior to sequencing enhances their similarity to unfractionated (W) samples.

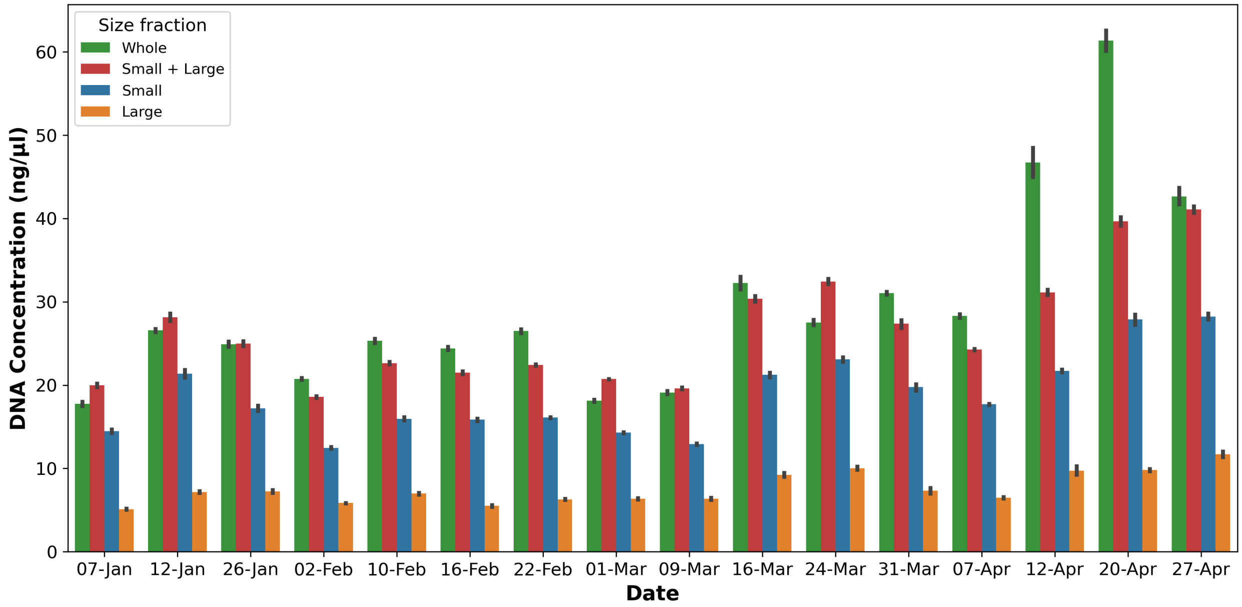


Figure 2: **DNA concentrations of size fractionated, unfractionated and de-fractionated samples over 16 sampling weeks**

Grouped bar plot of DNA concentration (ng/) of size fractionated (small (S), and large (L)), and unfractionated (W) samples from January 7, to April 27 2022 in the Bedford Basin. Each bar represents the DNA concentration from each size fraction at each given week. The data shows the differences in DNA yield, between size fractions and highlights the lower diversity observed in L in comparison to S, or W.

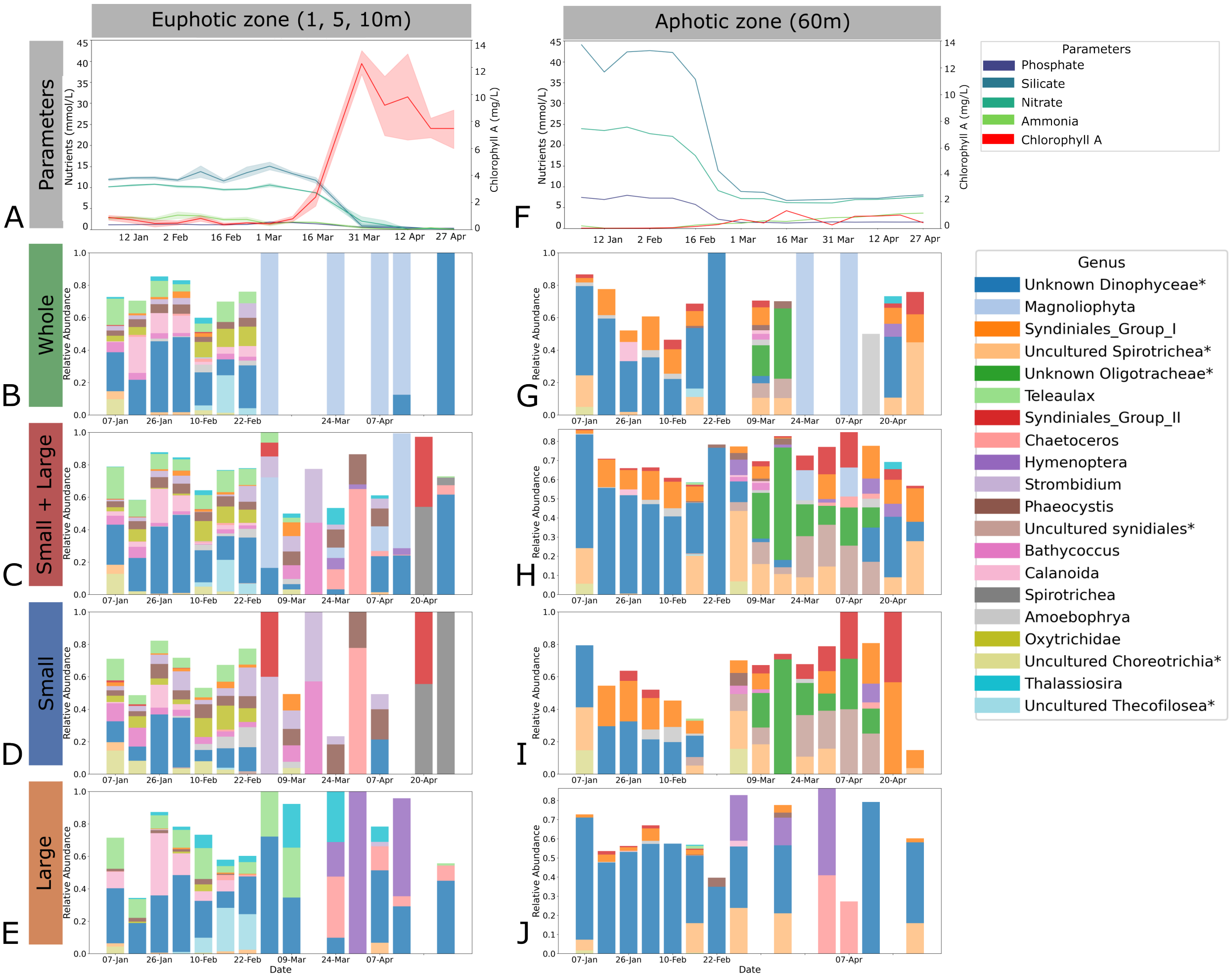


Figure 3: Concentrations of nutrients and chlorophyll a and eukaryotic microbial composition in the unfractionated (whole), de-fractionated (small + large) and fractionated (small, large) sets

Bar plots of the concentrations of nutrients and chlorophyll a in the euphotic (1m, 5m, and 10m) and aphotic zone (60m) measured from January 7 to April 27, 2022. Nutrients including phosphate, silicate, nitrate and ammonia are quantified in mmol/L and chlorophyll a in µg/L. In the euphotic zone, solid lines represent the mean values across depths 1m, 5m, and 10m and the shaded area indicates the 95% confidence intervals. Nutrient concentrations are represented by blue and green hues, and chlorophyll a in red. A secondary y-axis was used for chlorophyll a to account for different scales facilitating the visualization of its relationship with the nutrients. Below the nutrient plots, four taxonomic bar plots show the microbial composition for the whole (B, G), de-fractionated (C, H), small (D, I) and large (E,J) size fractions during the spring bloom depicting the 20 most abundant genera. Some abundant genera were labeled as unclassified or uncultured. To provide more clarity, they were reidentified as uncultured or unclassified, along with their closest available taxonomic rank (e.g., genus, family, or class, depending on availability). These labels are also marked with an asterisk. The 5m depth taxonomy was used for representing the euphotic zone.

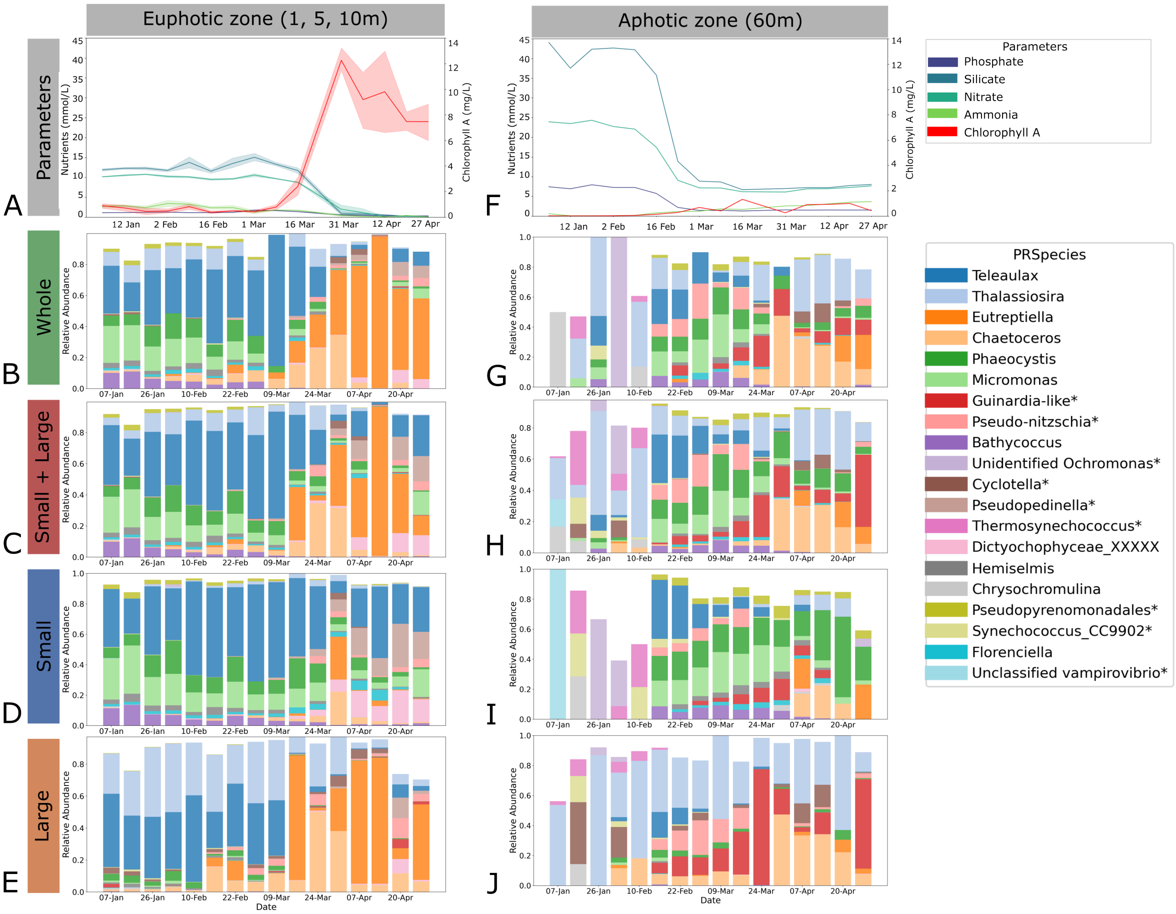


Figure 4: Chloroplast-associated taxonomy and environmental profile in the unfractionated (whole), defractionated (small + large) and fractionated (small, large) fractions

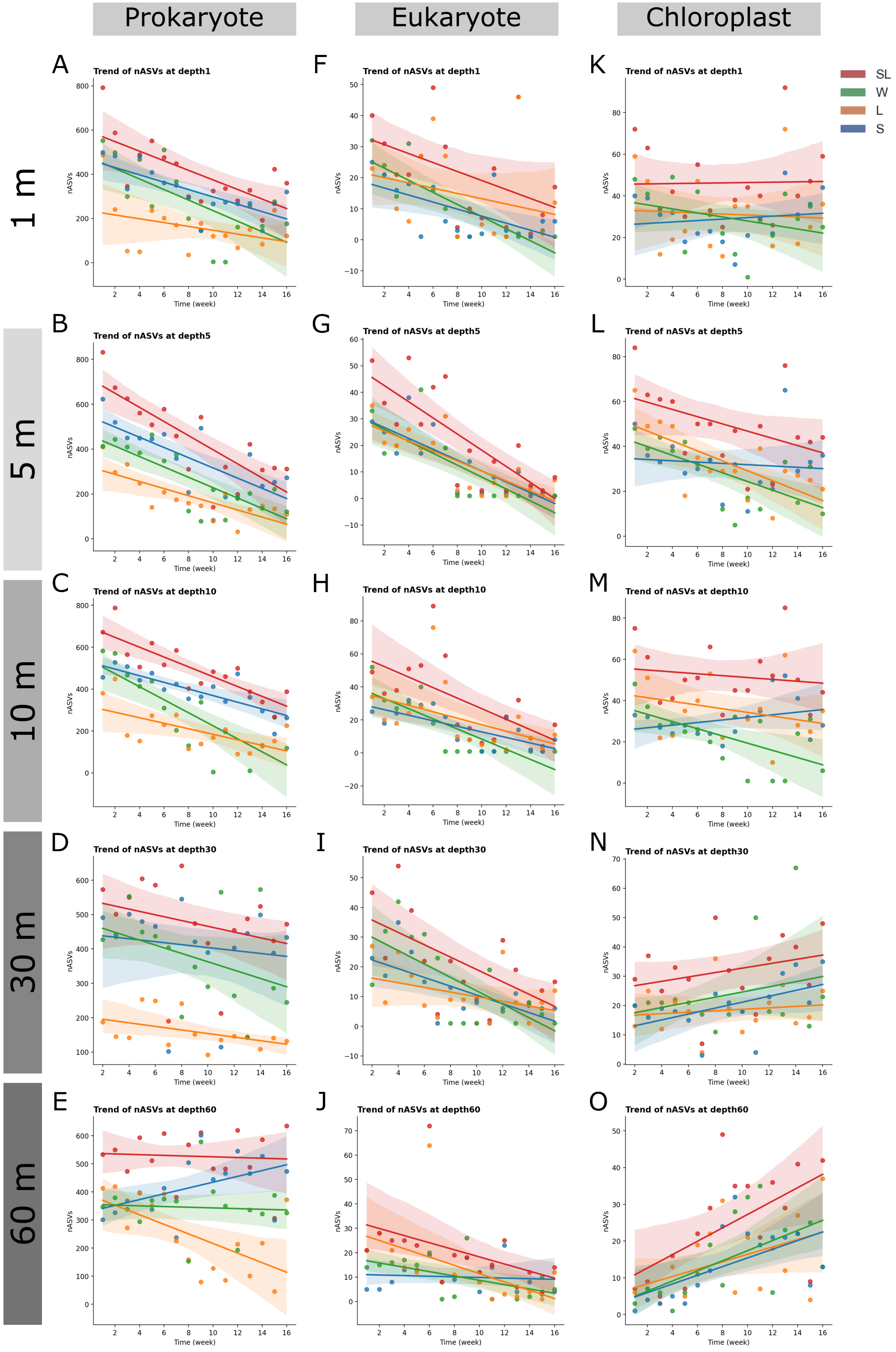


Figure 5: Trends

Linear regression of richness over time for each size fraction: large (L, orange), small (S, blue), unfractionated (W, green), and de-fractionated (SL, red) across depths 1, 5, 10, 30 and 60m. The panels are arranged by depth (rows), and community type (columns) prokaryotes, chloroplast and eukaryotes. The regression slopes assess whether the richness trends over the sample period are increasing, decreasing or stable.

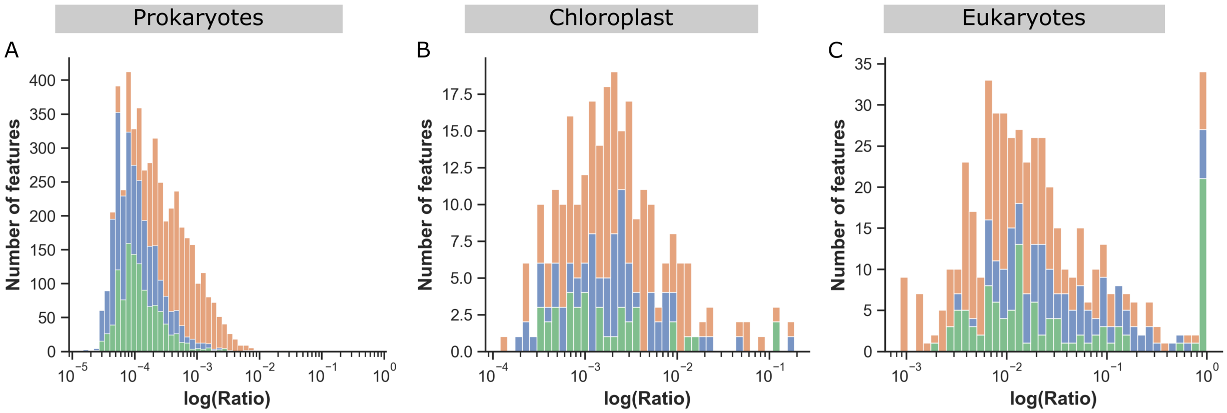


Figure 6: **Distribution of relative abundance of unshared features across size fractions**
Stacked distribution plots of unshared features for A) Prokaryotes, B) Chloroplast, C) eukaryotes categorized by size fraction.The right skewed distributions suggesting that most unshared features are rare, with some exception for the eukaryotes. In each bar, the total height represents the combined count of unshared small, large, and whole features and the size of the colored segments represent the size fraction contribution of that bin.

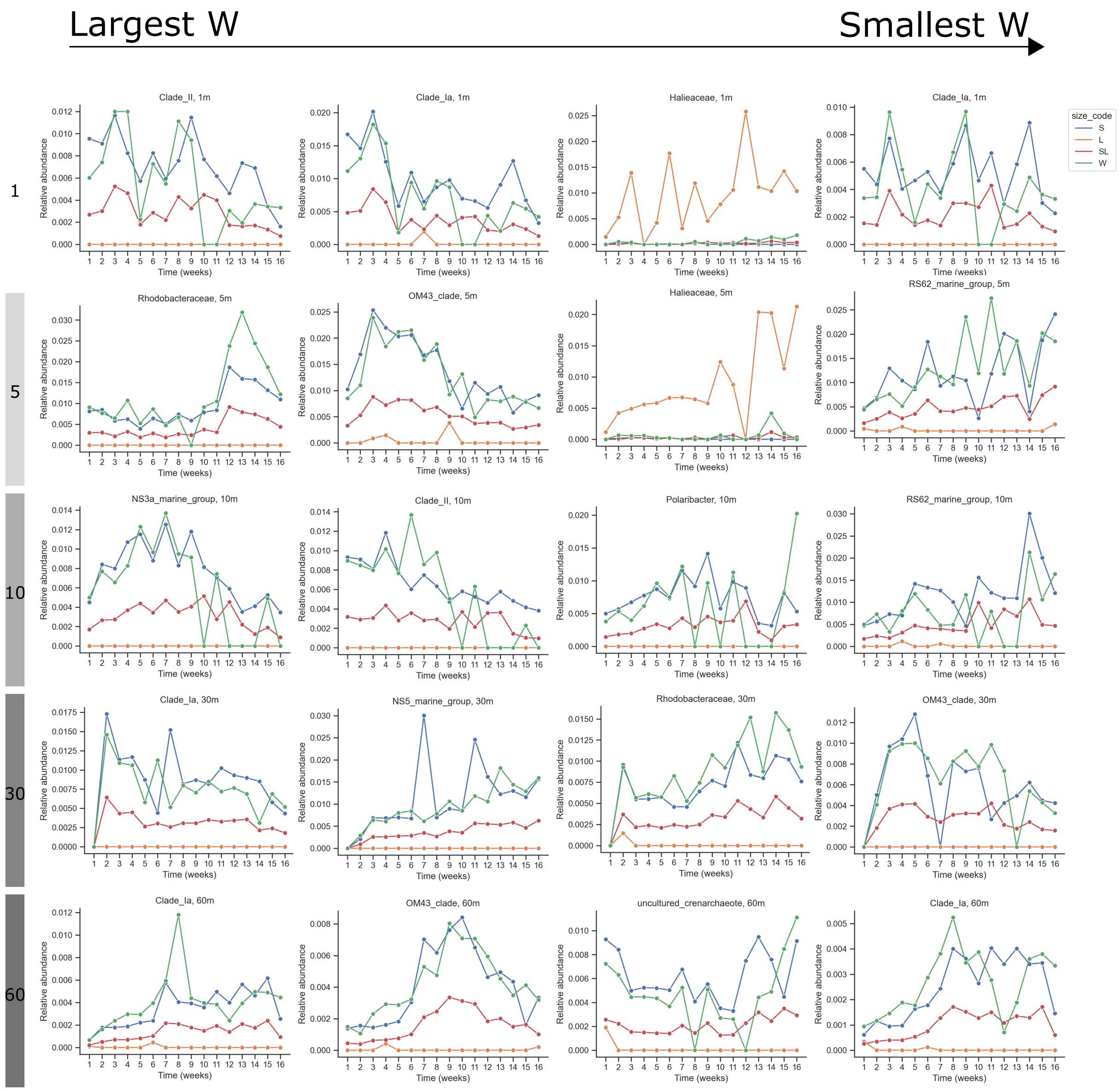


Figure 7: **Temporal dynamics of 12 features selected across filtration methods**
Time series of relative abundance for 12 sequence variants selected based on ANCOM raking with highest W values across depths, and taxonomic groups: prokaryotes (top row), eukaryotes (middle row), and chloroplast (bottom row). The time series span from January to April 2022, covering both pre-bloom and bloom periods. Each plot represents shows how relative abundance varies over time for samples separated among unfractionated, fractionated, and de-fractionated fractions. Differences in abundance patterns are observed both within and between taxa, highlighting the influence of size fractionation on community composition. These results suggest that the impact of size fractionation is taxon-specific and time-dependent.

| Depth | Pair | | Prokaryote | | | Chloroplast | | | Eukaryote | | |
|---|---|---|---|---|---|---|---|---|---|---|---|
| | | | R2 | p.adjusted | ANCOM | R2 | p.adjusted | ANCOM | R2 | Adj. p-value | ANCOM |
| 1 | W | L | 0.180 | **0.002** | 21 | 0.081 | **0.048** | 1 | 0.049 | 0.339 | 0 |
| | W | S | 0.043 | 0.216 | 0 | 0.056 | 0.110 | 0 | 0.026 | 0.937 | 0 |
| | W | SL | 0.034 | 0.380 | 1 | 0.024 | 0.694 | 0 | 0.022 | 0.944 | 0 |
| | L | S | 0.291 | **0.002** | 41 | 0.166 | **0.006** | 3 | 0.072 | 0.006 | 0 |
| | L | SL | 0.248 | **0.002** | – | 0.076 | **0.048** | – | 0.024 | 0.937 | – |
| | S | SL | 0.014 | 0.986 | – | 0.042 | 0.220 | – | 0.024 | 0.937 | – |
| 5 | W | L | 0.200 | **0.002** | 26 | 0.065 | **0.039** | 160* | 0.042 | 0.346 | 0 |
| | W | S | 0.053 | 0.092 | 1 | 0.075 | **0.038** | 159* | 0.039 | 0.377 | 0 |
| | W | SL | 0.043 | 0.156 | 0 | 0.038 | 0.249 | 0 | 0.019 | 0.991 | 0 |
| | L | S | 0.283 | **0.002** | 100 | 0.175 | **0.006** | 4 | 0.094 | **0.006** | 1 |
| | L | SL | 0.235 | **0.002** | – | 0.074 | **0.038** | – | 0.041 | 0.346 | – |
| | S | SL | 0.015 | 0.956 | – | 0.051 | 0.088 | – | 0.021 | 0.991 | – |
| 10 | W | L | 0.184 | **0.002** | 22 | 0.104 | **0.006** | 1 | 0.062 | 0.072 | 0 |
| | W | S | 0.079 | **0.012** | 3 | 0.087 | **0.011** | 1 | 0.052 | 0.100 | 0 |
| | W | SL | 0.064 | **0.032** | 0 | 0.066 | **0.036** | 0 | 0.038 | 0.290 | 0 |
| | L | S | 0.330 | **0.002** | **122** | 0.207 | **0.006** | 7 | 0.115 | **0.006** | 1 |
| | L | SL | 0.271 | **0.002** | – | 0.084 | **0.011** | – | 0.029 | 0.507 | – |
| | S | SL | 0.019 | 0.908 | – | **0.071** | **0.023** | – | 0.047 | 0.150 | – |
| 30 | W | L | 0.305 | **0.002** | 31 | 0.130 | **0.002** | 2 | 0.050 | 0.096 | 0 |
| | W | S | 0.044 | 0.224 | 0 | 0.069 | **0.026** | 0 | 0.039 | 0.285 | 0 |
| | W | SL | 0.034 | 0.524 | 0 | 0.037 | 0.308 | 0 | 0.033 | 0.590 | 0 |
| | L | S | 0.371 | **0.002** | 153 | 0.255 | **0.002** | 6 | 0.096 | **0.006** | 1 |
| | L | SL | 0.337 | **0.002** | – | 0.121 | **0.002** | – | 0.044 | 0.285 | – |
| | S | SL | 0.012 | 0.990 | – | 0.084 | **0.012** | – | 0.021 | 0.917 | – |
| 60 | W | L | 0.180 | **0.002** | 18 | 0.072 | **0.024** | 1 | 0.044 | 0.244 | 0 |
| | W | S | 0.038 | 0.350 | 0 | 0.066 | **0.038** | 1 | 0.053 | 0.096 | 0 |
| | W | SL | 0.032 | 0.434 | 1 | 0.032 | 0.377 | 0 | 0.030 | 0.671 | 0 |
| | L | S | 0.237 | **0.002** | 20 | 0.169 | **0.006** | 4 | 0.090 | **0.006** | 0 |
| | L | SL | 0.186 | **0.002** | – | 0.060 | **0.038** | – | 0.026 | 0.671 | – |
| | S | SL | 0.016 | 0.925 | – | 0.060 | 0.072 | – | 0.029 | 0.671 | – |

Table 2: Multi-depth statistical analyses: alpha diversity, and ANCOM results from prokaryotic, chloroplast and eukaryotic communities across size fractions

For each depth, and community type, pairwise comparisons among unfractionated (W), large (L), small (S), and de-fractionated (SL) sets are summarized. The table reports $R^2$ values, adjusted p-values, and ANCOM test results.

| **Depth** | **Category** | **Prokaryotes** | | Chloroplast | | Eukaryotes | |
|---|---|---|---|---|---|---|---|
| | | **parOmegaSq** | **Pr(>F)** | parOmegaSq | Pr(>F) | parOmegaSq | Pr(>F) |
| **1** | **Size fraction** | 0.152 | 0.001 | 0.068 | 0.001 | 0.006 | 0.25 |
| | **Time** | **0.016** | 0.033 | 0.020 | 0.024 | 0.021 | 0.02 |
| **5** | **Size fraction** | **0.158** | **0.001** | 0.075 | 0.001 | 0.016 | 0.094 |
| | **Time** | **0.012** | 0.060 | 0.018 | 0.025 | 0.009 | 0.087 |
| **10** | **Size fraction** | 0.192 | **0.001** | 0.110 | 0.001 | 0.039 | 0.001 |
| | **Time** | 0.015 | **0.047** | 0.012 | 0.054 | 0.015 | 0.034 |
| **30** | **Size fraction** | **0.220** | **0.001** | 0.126 | 0.001 | 0.019 | 0.018 |
| | **Time** | 0.013 | 0.074 | 0.028 | 0.003 | 0.007 | 0.092 |
| **60** | **Size fraction** | 0.129 | **0.001** | 0.073 | 0.001 | 0.017881 | 0.041 |
| | **Time** | **0.042** | **0.004** | 0.055 | 0.001 | 0.022850 | 0.012 |

Table 3: **Multi-depth PERMANOVA analysis of size fraction, and temporal effects on prokaryote, chloroplast and eukaryote microbial communities**

Summary of PERMANOVA results with reported partial omega squared (effect size), and p-value for assessing whether the microbial communities sampled in different size fractions, or at different time grouping (pre-bloom or bloom) differs. The partial omega squared value is the effect size, and the p-value is the significance of the hypothesis testing.
